# Similar levels of emotional contagion in male and female rats

**DOI:** 10.1101/857094

**Authors:** Yingying Han, Bo Sichterman, Maria Carrillo, Valeria Gazzola, Christian Keysers

## Abstract

Emotional contagion, the ability to feel what other individuals feel, is thought to be an important element of social life. In humans, emotional contagion has been shown to be stronger in women than men. Emotional contagion has been shown to exist also in rodents, and a growing number of studies explore the neural basis of emotional contagion in male rats and mice. These studies promise to shed light on the mechanisms that might go astray in psychiatric disorders characterized by dysfunctions of emotional contagion and empathy. Here we explore whether there are sex differences in emotional contagion in rats. We use an established paradigm in which a demonstrator rat receives footshocks while freezing is measured in both the demonstrator and an observer rat, which can hear, smell and see each other. By comparing pairs of male rats with pairs of female rats, we find (i) that female demonstrators freeze less when submitted to footshocks, but that (ii) the emotional contagion response, i.e. the degree of influence across the rats, does not depend on the sex of the rats. This was true whether emotional contagion was quantified based on the slope of a regression linking demonstrator and observer average freezing, or on Granger causality estimates of moment-to-moment freezing. The lack of sex differences in emotional contagion is compatible with an interpretation of emotional contagion as serving selfish danger detection.

## 1. Introduction

Affective empathy, i.e. feeling what another feels while knowing that the other person’s affective state is the source of our own affective state ^1,2^, has often been reported to show gender differences in humans ^3–5^, with women more affected by the emotions of others. Many believe that empathy evolved in the context of parental care, where feeling the distress of offspring motivates nurturing behavior and thereby increases Darwin fitness ^2,6,7^. If empathy serves maternal care, one may predict empathy to be stronger in females.

Emotional contagion is thought of as an evolutionary predecessor to empathy ^6–9^. The term emotional contagion can be traced back to the German ‘Stimmungsuebertragung’ introduced by Konrad Lorentz to refer to cases in which witnessing a conspecific in a particular emotion, expressed via movements and sounds, triggers a similar emotion in the witness (“der Anblick des Artgenossen in bestimmten Stimmungen, die sich durch Ausdrucksbewegungen und -laute äußern können, im Vogel selbst eine ähnliche Stimmung hervorruft”^10^). Similar to empathy, in humans, there is some evidence that emotional contagion is more pronounced in women ^4,11–13^, and female babies are more likely to cry and cry for longer when hearing another baby cry ^14,15^.

Mounting experimental evidence suggests that rats and mice show signs of emotional contagion ^16–23^. Would they also show sex differences in emotional contagion with females showing more contagion than males? Emotional contagion can be quantified systematically in rodents using designs in which one demonstrator animal receives a footshock, and the freezing of another observer that witnesses the event is found to be increased, suggesting that the distress of the shocked demonstrator was transferred to the observer ^16,17,21,24–29^. We have recently introduced ways to quantify emotional transfer in this paradigm by leveraging Bayesian statistics and Granger causality ^29^. Here we therefore ask whether by using these quantification methods, emotional contagion is stronger in females compared to male Long Evans rats. With mounting interests in the biological and neural basis of emotional contagion ^21,22,26,28,30,31^, exploring sex differences is important. This is particularly true in a paradigm involving the nociceptive system and freezing as behavioral read-out because profound sex differences exist in the biology of the nociceptive system ^32^, and female rats have been shown to be generally more active than males and respond to shocks with less freezing ^33,34^.

To date the majority of studies investigating sex differences on emotional contagion have been conducted in mice ^23,25,27,35–39^. In fear observation paradigms, in which the response of an observer mouse is measured while it witnesses another experiencing a negative stimulus (generally a footshock), evidence from mice is contradictory: while Keum et al. (2016), Sanders et al. (2013) and Chen et al. (2009) find no effect of sex, Pisansky et al., 2017 finds that 1) females mice have a greater amount of emotional contagion as measured by higher amounts of freezing, 2) in contrast to males, familiarity does not play a role in female mice, and 3) this response is modulated by oxytocin. These opposing effects between studies could be due to differences in the shocking protocols (e.g., a much longer testing session in Pisansky compared to the other three studies) and the freezing quantification method. In paradigms measuring emotional contagion through pain hypersensitivity, female mice display a larger amount of socially transferred pain hypersensitivity compared to males ^38^. Noteworthy, in these type of paradigms, the same pattern of the familiarity effect observed in Pisansky has been reported ^23,36^: while female mice have an equivalent response when with a familiar and an unfamiliar animal, males show a reduced effect when tested with an unfamiliar conspecific. However, this effect is reversed in paradigms measuring approach to conspecifics in distress ^35^: female mice approach distressed cagemates more than unfamiliar animals, and males do not differentiate between familiar vs unfamiliar demonstrators. This approach effect seems to be conserved across species as the same phenomenon is observed in rats ^40^.

Overall, in rats there is a much smaller number of studies that have investigated the effect of sex on emotional contagion, with no studies that have measured the effect of sex in fear observation paradigms, inconclusive results in a study investigating social buffering (sex effects that go in opposite direction at the behavior and endocrine level ^41^) and one study that finds subtle sex differences on two-way avoidance learning (an indirect proxy measure for emotional contagion ^42^). Lastly, only one study n rodents^43^ has reported sex effects in a paradigm measuring prosociality in rats, where they find that females are more likely to help a conspecific in distress than males, supporting the notion that females are more empathic than males. The sparsity of studies in rats and the non-converging results in mice motivated us to apply our new analytical approach to investigate sex differences in emotional contagion in rats.

To compare emotional contagion in males and female rats, we thus harness a paradigm developed in our lab in which a shock-experienced observer rat interacts through a perforated transparent divider with a demonstrator rat receiving footshocks. We quantify the freezing behavior of both animals during an initial 12-minute baseline period and a 12-minute test period in which the demonstrator received 5 footshocks (1.5mA, 1s each, ISI: 120 or 180s, Fig. 1). We have previously shown that there is *mutual* influence across demonstrators and observers, with the distress of the shocked demonstrators triggering freezing in observers and individual differences in observer reaction influencing back how much demonstrators freeze (Han et al., in press). In addition to (i) comparing the average freezing level of observers across the two sexes, here we therefore additionally (ii) compare emotional contagion in terms of the *relationship* between the freezing of observers and demonstrators in terms of average freezing during the shock period using Bayesian regression analyses and (iii) in terms of second-to-second freezing influences using Granger causality (Han et al., in press).

**Figure 1.**
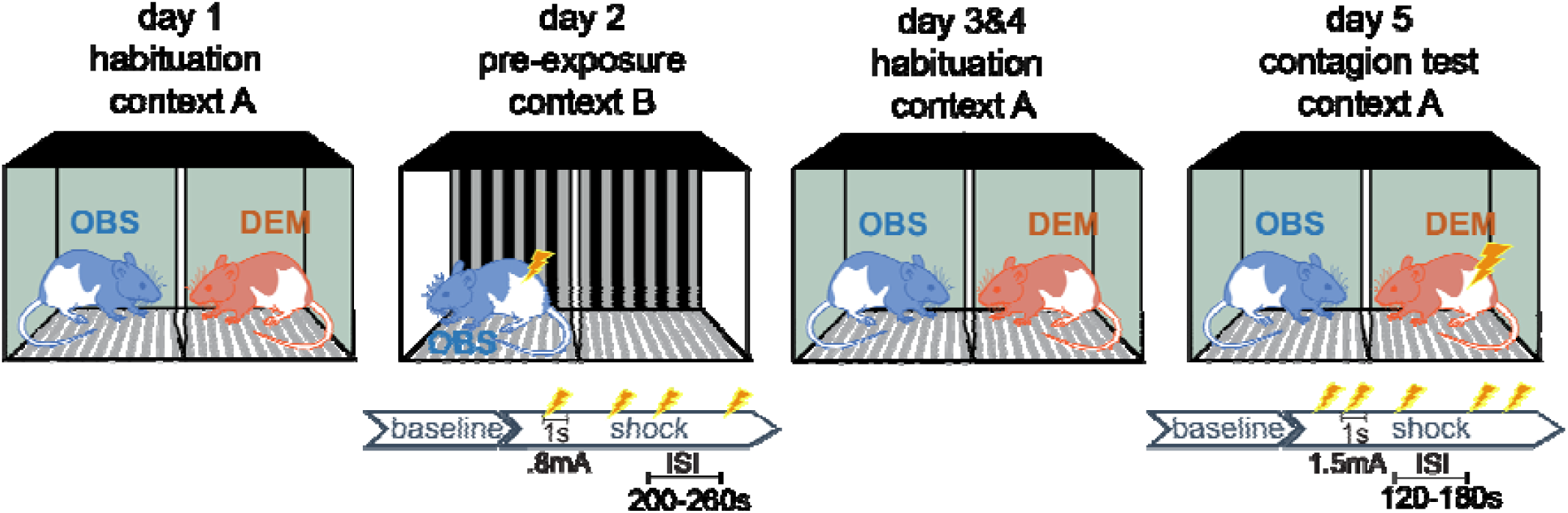
Timeline of the emotional contagion test. Following the first day of habituation, in the shock pre-exposure session, the observer animals were exposed to footshocks alone in a context that is different from the test apparatus (day 2). The pre-exposure session was followed by two more days of habituation (day 3 & 4) to reduce contextual fear in the contagion test session. On day 5, demonstrator-observer dyads were placed in the setup for a total of 24 minutes. After a 12 minutes baseline period which is identical to the habituation session, the demonstrators received 5 footshocks (each 1.5mA, 1sec long) during the 12 minutes shock period. The inter-shock intervals were either 2 or 3 minutes.

Given that in the past we had run an experiment with only females ^16^, and later several experiments with only males ^17,28,29^, and observed higher levels of freezing in both male observers and male demonstrators than in our original female sample, we expected that females would show reduced levels of freezing compared to males in this experiment. However, when conceiving of emotional contagion as the link between the freezing of the two animals, would we find a stronger link in females than males?

## 2. Results

### 2.1. Average freezing is higher in male rats

Figure 2A shows the group freezing data of observers and demonstrators of both sexes. A test of normality revealed that freezing during the shock epoch is normally distributed for both sexes and roles (Shapiro-Wilk, all *p*>0.3). Unfortunately, freezing during baseline deviates from normality for both sexes and roles (Shapiro-Wilk, all *p*<0.05). Accordingly, parametric tests including the baseline should be interpreted with caution, and will be supplemented by non-parametric tests where possible.

**Figure 2:**
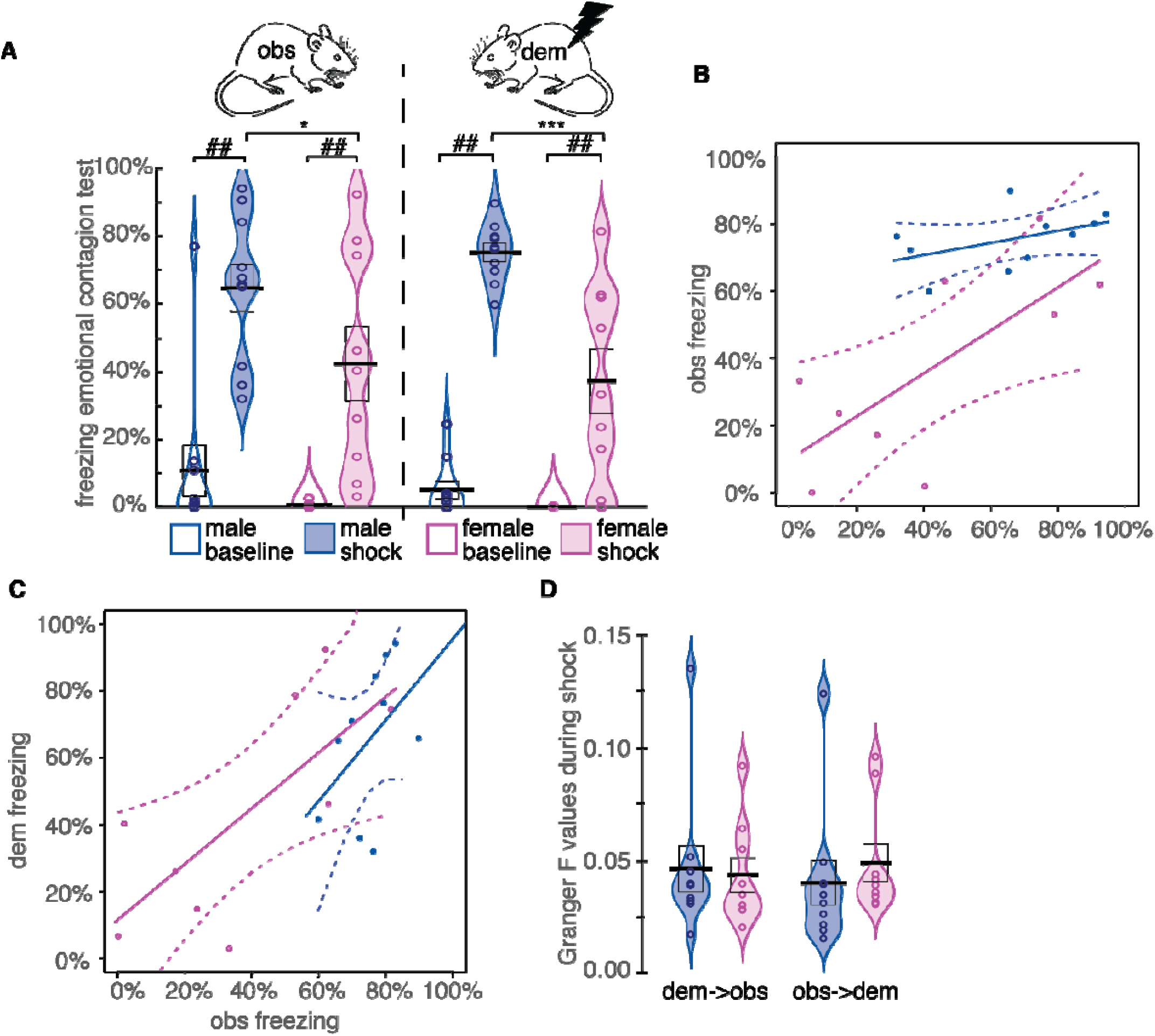
Emotional Contagion as a function of sex. (A) Freezing percent during the baseline (open violins) and the shock period (filled violins) for male (blue) and female (purple) rats, with observer data on the left and demonstrator data on the right. The black bar represents the mean, the box ± sem. (B) Observer freezing as a function of demonstrator freezing including linear regression lines and their 95% confidence intervals. (C) Demonstrator freezing during the shock period as a function of observer freezing during the shock period including linear regression lines and their 95% confidence intervals. (D) Granger causality *F* values in the dem->obs (left) and obs->dem (right) direction. (B-D) only present freezing from the shock epoch. For all panels: *:*p*<0.05, **:*p*<0.01, ***:*p*<0.001, two-tailed *t*-test ##=*p*<0.005 in Wilcoxon test. Other conventions are the same as in (A).

For observers a 2 Sex (male vs female) x 2 Epoch (baseline vs shock), revealed a main effect of Epoch (*F*(1,17)=56, *p*<0.001, BF_incl_=874958), a trend for Sex (*F*(1,17)=3.86, *p*=0.066, BF_incl_=1.3) but no interaction (*F*(1,17)=1.002, *p*=0.33, BF_incl_=1.3). Paired tests comparing observers’ freezing during the baseline and the shock period confirm that witnessing the demonstrator receive shocks increases freezing compared to baseline in both sexes examined individually (Wilcoxon *W*(9) = 0, *p*=0.002 for males; *W*(8) = 0, *p=*0.004 for females; Fig. 2A). We then compared the freezing of the male versus the female observers during the shock period. Considering the findings from previous research, we had hypothesized lower freezing in females compared to male observers. A one-tailed *t*-test on the observer freezing confirmed this prediction (*t*(17)=1.8, *p*<0.044), with a large effect size with males freezing 1.5 times as much as females (Cohen *d*=0.8, Fig. 2A).

For demonstrators a 2 Sex (male vs female) x 2 Epoch (baseline vs shock) on freezing also revealed a main effect of Epoch (*F*(1,17)=115, *p*<0.001, BF_incl_=1E10), but also revealed a significant main effect for Sex (*F*(1,17)=19, *p*<0.001, BF_incl_=208) and an interaction (*F*(1,17)=11, *p*=0.004, BF_incl_=103), due to a larger increase in freezing following the shocks in males. A one-tailed *t*-test again confirms that during the shock epoch, male demonstrators froze more than female demonstrators (*t*(17)=3.97, *p*_1_<0.001) with an even larger effect size than for the observers: males froze twice as much as females (Cohen *d*=1.8, Fig. 2A).

For both observers and demonstrators, males thus froze more than females. That the effect size was more pronounced for demonstrators (*d*=1.8) than observers (*d*=0.8) raises the questions of whether the smaller sex difference in observers is a downstream result of the larger sex difference in demonstrators, a question we will address in the next section.

### 2.2. Demonstrator freezing level, independently of sex, is the best predictor of observer freezing

To understand whether sex differences in observers’ freezing during the shock epoch (obs_f_) were due to differences in the demonstrators’ freezing (dem_f_) during the shock period alone we performed an ANCOVA, which tests whether after regressing out the individual differences in demonstrator freezing (dem_f_), there is a residual main effect of observer sex, or an interaction of dem_f_ and sex. This was done using a frequentist and a Bayesian ANCOVA with 2 sex (male vs female) x dem_f_, see Table 1A.

**Table 1:**
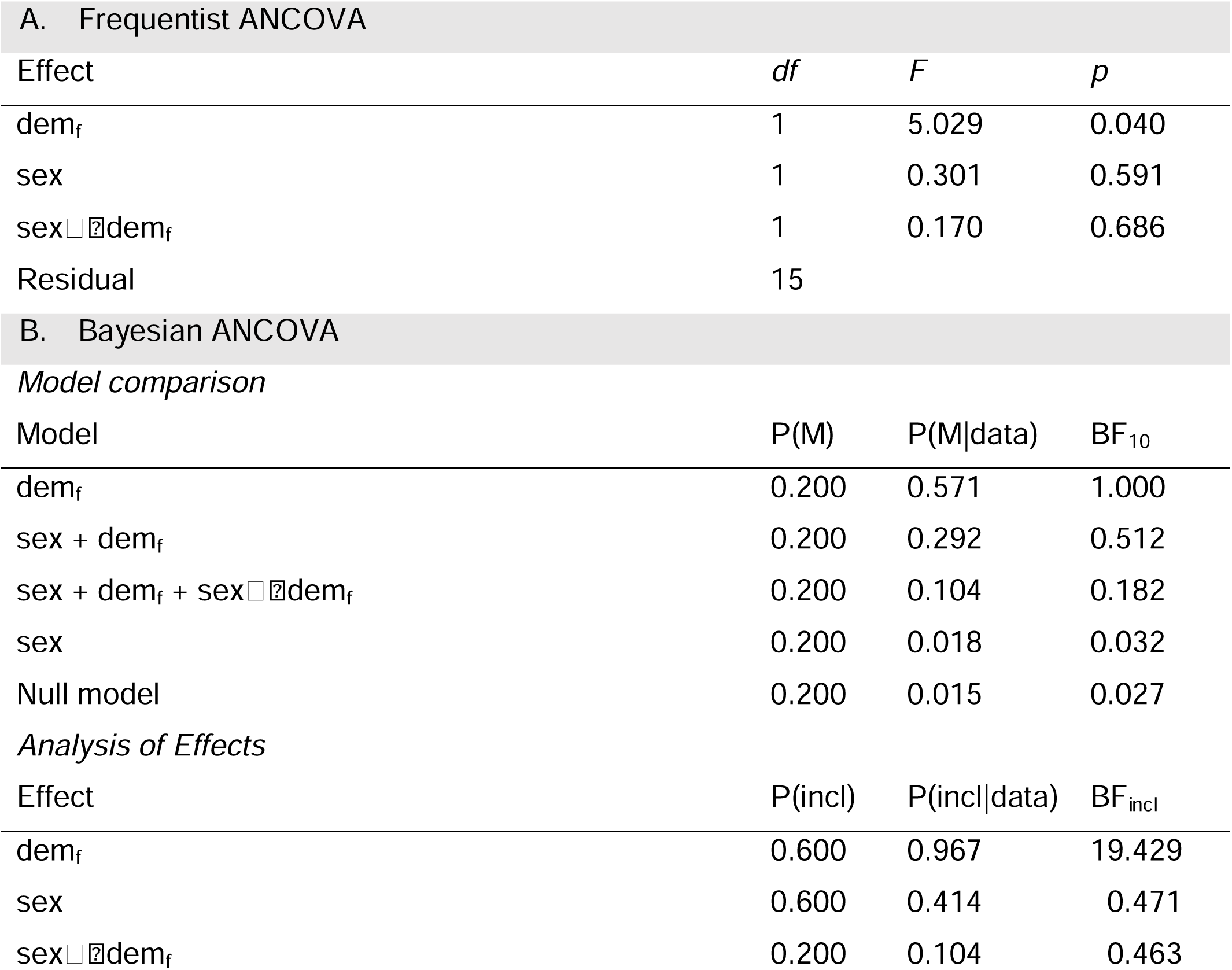
Analysis of observer freezing considering the shock period only. Observer freezing was analyzed using demonstrator freezing (dem_f_), sex (1=male, 0=female), and their interaction as explanatory variables. Null models only include an intercept. The Bayesian ANCOVA was performed in JASP using default priors (sex=fixed factor, dem_f_=covariate, prior *r* on sex=0.5 on dem_f_=0.354), and models are ranked based on their predictive credibility. For the model comparison, P(M) refers to the prior likelihood of each model, P(M|data) the posterior likelihood of the model given the observed data. BF_10_ quantifies the relative evidence of the models compared to the best model. For the analysis of effects, P(incl) refers to the prior likelihood of including an effect in the model, P(incl|data) the posterior likelihood after having seen the data, and BF_incl_ is the BayesFactor for inclusion of an effect, i.e. the likelihood of the data under models including the factor divided by that of models not including the factor.

The frequentist analysis revealed a significant influence of dem_f_ confirming that the observer freezing reflects the demonstrator freezing but neither a significant main effect of sex or interaction of sex * dem_f_ were detected. This shows, that once the difference in demonstrator freezing have been accounted for, sex fails to explain additional variance. This suggests, that difference in observer freezing can be most parsimoniously explained by knowing the level of freezing of the demonstrator, independently of sex. In Fig. 2B, this is apparent in the linear regression lines that are very similar across the two sexes, with clearly overlapping confidence intervals, suggesting that the differences in demonstrator freezing simply acted as distinct input onto a transmission function (i.e. slope and offset) that is the same independently of the sex of the observer. Given that female demonstrators reacted to the shocks with less freezing, this simply transforms into the group difference in observer freezing we observe.

A non-significant main effect of sex or interaction with sex could reflect evidence that there is no effect of sex (evidence of absence), or that our study was underpowered and cannot speak for or against the absence of a sex difference. To shed light on this issue, we performed a Bayesian ANCOVA that explains observer freezing using competing models with or without sex as a factor (Table 1B). A model only considering dem_f_ is the best model by a margin (P(M|data), Table 1B), and an analysis of effects provides strong evidence for an effect of dem_f_ (BF_incl_=19), confirming that the demonstrator freezing strongly determines observer freezing. The analysis also shows that the evidence leans towards the *absence* of an effect of sex, be it as a main effect (BF_incl_=0.471) or sex*dem_f_ interaction (BF_incl_=0.463), showing that the data is over twice as likely in models without these factors than in models with them. Indeed, a full model in which sex and its interaction are included is 5 times less likely than one only including dem_f_. Altogether this shows that our data is best explained by a model that considers the level of freezing of the demonstrator (which is different for male and female demonstrators, as shown above) but ignores the sex of the animals involved.

To further characterize the relation between observers’ and demonstrators’ freezing in the two sexes, we performed separate Bayesian regressions for males and females. This gave highly overlapping posterior estimates for the regression weight for obs_f_ = *β*\*dem_f_ + intercept, with *β* for females having mean=0.44 (95% credibility interval CI=[0.0,1.09]) and male=0.36(95%CI=[-0.5,1.9]). Fig. 2B illustrates this as the similarity in slope.

In our past work, we have shown that differences in the freezing level of the observer can influence back the freezing level of the demonstrator, a phenomenon akin to social buffering. To explore whether there might be a sex difference in this influence of obs_f_ on dem_f_, we performed a frequentist ANCOVA and a Bayesian model comparison between different models explaining freezing of the demonstrators as a function of sex, freezing of observers (obs_f_) and their interaction (sex*obs_f_, Table 2). The frequentist ANCOVA showed significant main effects and a trend for an interaction (*p*=0.129, Table 2A). The Bayesian model comparison found that including sex and observers’ freezing in an additive model (obs_f_ + sex) best describes the data (Table 2B). There was strong evidence for a contribution of sex in predicting demonstrator freezing (BF_incl_=14.712), with the females freezing less to the shock than the males. There was also strong evidence for a contribution of observer freezing (BF_incl_=18.390). However, there was only a trend and anecdotal evidence for including an interaction effect (BF_incl_=2.804) indicating that if there was a sex difference in social buffering, we would need a larger group to find robust evidence for such an effect. This apparent difference in slope across the sexes can be appreciated in Fig. 2C.

**Table 2:** Analysis of demonstrator freezing. Demonstrator freezing was analyzed using observer freezing (obs_f_), sex (1=male, 0=female), and their interaction as explanatory variables. Null models only include an intercept.

| A. Frequentist ANCOVA |  |  |  |
| --- | --- | --- | --- |
| Effect | *df* | *F* | *p* |
| $obs_f$ | 1 | 8.2 | 0.012 |
| sex | 1 | 8.5 | 0.011 |
| $sex \square \square obs_f$ | 1 | 2.6 | 0.129 |
| Residual | 15 |  |  |
| B. Bayesian ANCOVA |  |  |  |
| Model comparison |  |  |  |
| Models | P(M) | P(M\|data) | $BF_{10}$ |
| $obs_f + sex$ | 0.200 | 0.511 | 1.000 |
| $obs_f + sex + obs_f \square sex$ | 0.200 | 0.412 | 0.807 |

| obs<sub>f</sub> | 0.200 | 0.042 | 0.083 |
| --- | --- | --- | --- |
| sex | 0.200 | 0.034 | 0.066 |
| Null model | 0.200 | 0.001 | 0.002 |
| *Analysis of Effects* |  |  |  |
| Effect | P(incl) | P(incl\|data) | BF<sub>incl</sub> |
| obs<sub>f</sub> | 0.600 | 0.965 | 18.390 |
| sex | 0.600 | 0.957 | 14.712 |
| obs<sub>f</sub>×sex | 0.200 | 0.412 | 2.804 |

### 2.3. No sex difference in Granger-causality across the animals

To further explore whether males and females differ in the temporal coupling of the freezing between demonstrators and observers, we performed Granger causality analyses (Fig. 2D). Unlike the other analyses that explore the average freezing over the 12 minutes of the shock period, the Granger causality analyses explore the relation between the second-to-second freezing of demonstrators and observers. Specifically, it examines if past demonstrator freezing can explain present observer freezing (to quantify influences in the dem→obs direction), and if past observer freezing can explain present demonstrator freezing (to quantify influences in the obs→dem direction). Higher G-causality values (i.e. Granger *F* values) indicate higher temporal coupling of the behavior of the two animals in a pair, and thereby stronger social sensitivity to the behavior of the other. A Granger analysis considering all animals (irrespective of sex) revealed significant information flow in both directions (dem→obs Granger *F* = 0.039, *p* < 0.0001; obs→dem Granger *F* = 0.034, *p* < 0.0001). Because the G-causality values were not normally distributed (Shapiro *p*<0.05), we use non-parametric tests to compare the sexes. We found no significant difference in Granger causality in either direction (Mann-Whitney *U*, dem→obs, *U*(17)=38, *p2*=0.6; obs→dem, *U*(17)=41, *p*_*2*_=0.78; Fig. 2D). Bayesian Mann-Whitney *U* tests revealed that in both directions, the evidence leans towards the null hypothesis, but with limited evidence strength: in dem→obs direction, the Bayes factor in favor of the null hypothesis BF_01_= 2.4, in the obs→dem direction BF_01_=2.1.

### 2.4. A trade-off between rearing and freezing

The finding that the female demonstrators froze less than their male counterparts raises the question of whether they reacted to the shocks using an alternative strategy. It has often been described that individual and sex differences exist in the propensity to react to danger with escape vs freezing ^33^. We thus explored whether females reared (including attempts to climb out of the box) more than their male counterparts (Fig. 3A). All rearing data were normally distributed (Shapiro *p*>0.1) except for male observer rearing during the shock epoch.

**Figure 3:**
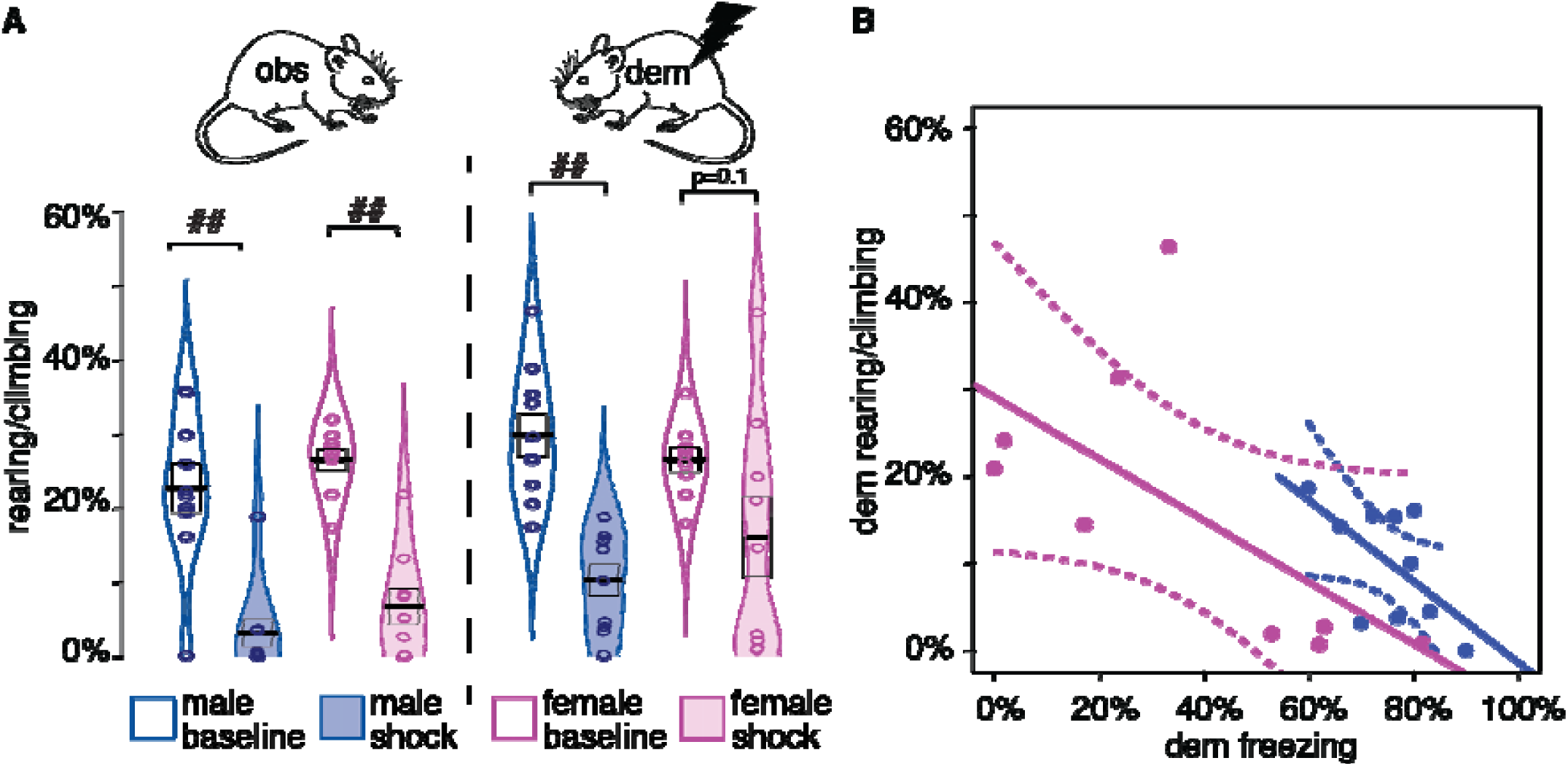
Rearing/climbing as a function of sex. (A) distribution of rearing and climbing. (B) The trade-off of rearing and climbing during the shock epoch for demonstrators. All conventions as in Figure 2. ##=*p*<0.01 in the Wilcoxon test.

All groups showed reduced rearing/climbing following the shocks. To explore if that reduction is sex-dependent, we performed mixed frequentist ANOVAs separately for observer rearing and demonstrator rearing, including 2 Sexes (male vs female) x 2 Epochs (baseline vs shock). We found main effects of Epoch in both cases (Obs: *F*(1,17)=67, *p*<0.001, BF_incl_=8.7E7; Dem: *F*(1,17)=25, *p*<0.001, BF_incl_=2478), but no main effect of Sex (Obs: *F*(1,17)=2.4, *p*=0.141, BF_incl_=0.66; Dem: *F*(1,17)=0.121, *p*=0.732, BF_incl_=0.56) or interaction of Sex x Epoch (Obs: *F*(1,17)=0.004, *p*=0.95, BF_incl_=0.66; Dem: *F*(1,17)=2.3, *p*=0.146, BF_incl_=1.25).

We also observed a consistent negative correlation between rearing and freezing in our animals (Fig. 3B). To explore that relationship further, we performed ANCOVAs that explore rearing during the shock period as a function of sex, freezing and sex * freezing, separately for observers and demonstrators. In both cases, the effect of freezing was negative (Obs: *F(1,15)*=11.8, *p*=0.004, BF_incl_=21.5; Dem: *F(1,15)*=4.15, *p*=0.06, BF_incl_=6.07) while the effect of sex (Obs: *F*(1,15)=0.12, *p*=0.73, BF_incl_=0.39, Dem: *F*(1,15)=0.30, *p*=0.59, BF_incl_=0.5) or sex*freezing interaction (Obs: *F*(1,15)=0.13, *p*=0.72, BF_incl_=0.4, Dem: *F*(1,15)=0.08, *p*=0.78, BF_incl_=0.55) were negligible, suggesting a sex-independent tradeoff: the more an individual freezes, the less it rears, and vice-versa, confirming the notion that animals that froze less reared more. These alternative allocations are particularly visible amongst female demonstrators (Fig. 3B). While most male demonstrators consistently showed high levels of freezing (>50%) and low rearing/climbing (<20%) during the shock period, amongst the females, about half showed a similar pattern whilst the other half showed a pattern only seen in females, with lower levels or freezing (<50%) but higher levels of rearing/climbing (>20%).

### 2.5. No sex differences in freezing to shocks during pre-exposure freezing/rearing

Considering the large sex differences in demonstrator freezing in response to shocks, we also analyzed freezing levels during pre-exposure, where the observers experience shocks (Fig. 4A). Based on the results from the demonstrator rats, we expected male observers to show about twice as much freezing as female observers. Because freezing in the males during pre-exposure was not normally distributed (Shapiro, *W*=0.7, *p*<0.001), we used non-parametric tests. Tests revealed significant increases in freezing from baseline to shock in both sexes (Wilcoxon, female: *W*=0, *p*<0.004, males: *W*=0, *p*<0.002). While during baseline, males showed significantly higher freezing levels in response to a novel environment (Mann-Whitney *U*=13, *p*=0.008), no sex-driven differences in freezing were detected during the shock epoch (Mann-Whitney *U*=39, *p*=0.66). Females during the shock period of this pre-exposure froze much more (*mean* = 92%; *SEM* = 3%), than their female demonstrators later did in the contagion test (mean = 37%; SEM = 29%, Mann-Whitney U=0, p<0.001).

**Figure 4:**
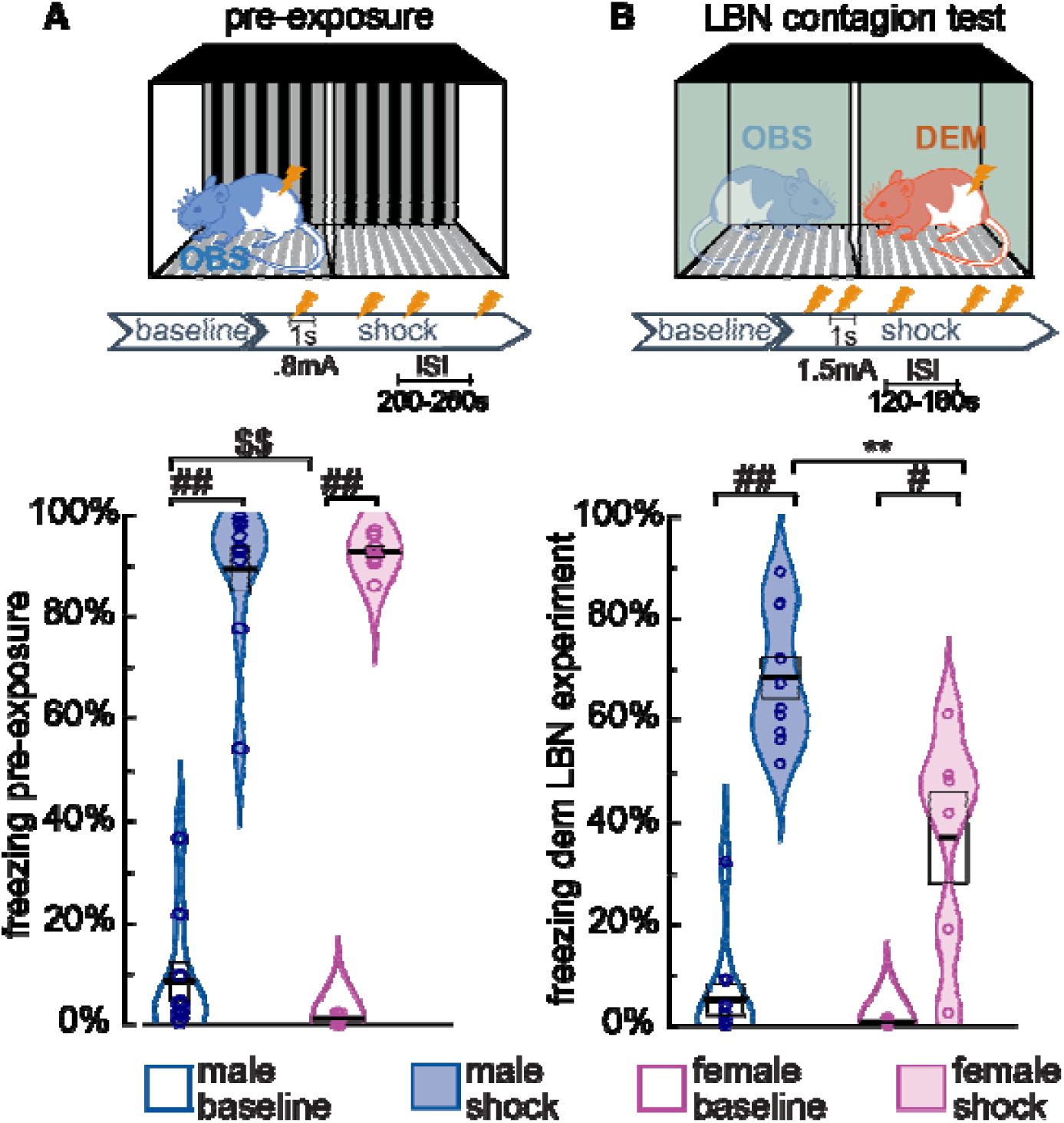
Freezing level for animals receiving shocks. (A) freezing in observer animals receiving shocks during pre-exposure. (B) freezing of demonstrators receiving shocks during the contagion test of the limited bedding and nesting (LBN) pilot group. The experimental schema above the panels illustrates the shock parameters and the fact that animals were alone in a new context in (A) but together with another animal in a familiar context in (B). ##: Wilcoxon test, *p*<0.01, #: Wilcoxon test, *p*<0.05. $$: Mann-Whitney *U, p*<0.01. **: *t*-test *p*<0.01.

To explore whether observers that froze more during pre-exposure also froze more while observing their demonstrator receive shocks, despite the non-normality of the male pre-exposure freezing, we tentatively performed an ANCOVA on observer freezing during the shock epoch that included sex as a fixed factor and demonstrator freezing and observer freezing during pre-exposure as covariates. The analysis confirmed that demonstrator freezing explains observer freezing during the shock epoch (*F*(1,15)=8.3, *p*=0.011, BF_incl_=22), but neither sex (*F*(1,15)=0.08, *p*=0.78, BF_incl_=0.5) nor pre-exposure freezing do (*F*(1,15)=1.23, *p*=0.28, BF_incl_=0.6).

Finally, the discrepancy between the strong sex effect found in demonstrators during the test situation (Fig. 2A) and its apparent absence during pre-exposure (Fig. 4A) raises the question of whether the sex-difference during the contagion test was simply a false positive. To explore this possibility, we drew on data from another group of animals in our experiment. Observer animals in that second group had undergone a limited bedding and nesting (LBN) manipulation as pups and are thus not reported here because they might not be representative of normal behavior. However, demonstrators in that LBN experiment were not submitted to limited bedding and were purchased in adulthood together with those of the main experiment and instead randomly assigned to the control group (that is reported in the method section in this paper). This LBN group can thus be used to confirm the presence or absence of a sex difference in response to shocks in the contagion test. The LBN demonstrators showed the same effect as the demonstrators in our main group (Fig. 4B). Freezing data was normally distributed during the shock period (Shapiro *p*>0.4), but not the baseline period (Shapiro *p*<0.01). During the shock epoch, male LBN demonstrators froze about twice as much as the females during the shock epoch (males: *n*=10, mean=68%, sem=4%; females *n*=6, mean=36%, sem=9%, *t*(14)=3.6, *p*=0.003, Cohen’s *d*=1.9, BF_10_=13), just as in the main experiment. While this LBN group might still be affected by slightly atypical observer reactions, it confirms that the sex difference found in our main group is robust. To explore whether the difference in freezing between pre-exposure and test could be driven by differences in the shocking protocol (pre-exposure: longer inter-shock intervals and fewer shocks), we also looked at the first 110s after the first 3 shocks in both sessions. Results showed qualitatively similar results, with significant sex differences in the contagion but not the pre-exposure freezing.

## Discussion

Based on the literature showing less freezing in females in non-social experiments ^33,34^, and our own experiments showing lower levels of freezing in female observers and demonstrators in our emotional contagion tests compared to males (see ^16^ for females and ^17,28,29^ for females), we wanted to directly test the effect of sex on our emotional contagion test, and expected to find reduced freezing in females compared to males. This expectation was confirmed by our data. However, the specific question of interest is whether there were sex differences in emotional contagion, the degree of affective alignment between two rats. Following up on our recent introduction of dyadic methods, that quantify the relationship between freezing behavior in observers and demonstrators ^29^, here we leverage these methods to quantify emotional contagion as the strength of the link between the freezing behavior across the two members of each pair. We exploit two methods to do so.

First, we used a regression analysis, which explores the relationship between average freezing of demonstrators and observers during the shock epoch, and found that a model that assumes the same slope for males and females is in fact the best description of our data. That is to say, that male and female observers react with the same amount of freezing to a given degree of freezing of the demonstrator. The sex differences we see in the overall freezing level of observers are thus not due to a difference in emotional contagion (i.e. a difference in sensitivity to demonstrator freezing) per se, but a result of a difference in the amount of freezing displayed by the demonstrator. This is particularly visible in Fig. 2B: female observers paired with those female demonstrators that displayed male-typical levels of freezing (>50%) showed a male-typical level of freezing (>50%).

Second, we used Granger causality to explore the moment-to-moment relationship between the freezing of the members of each pair. We found reliable, bidirectional evidence of influence across the animals in both male and female dyads, and there was no reliable difference in the strength of this Granger causality. This shows again, that the social transfer of distress, as measured by the relationship in freezing behavior across the members of the dyads was independent of sex.

Overall, because our study had a relatively small sample size (10 male and 9 female dyads), we also consistently used Bayesian statistics to explore whether a lack of significant sex-difference in our frequentist approaches was simply due to a lack of power (and thus represents absence of evidence) or provided evidence for the lack of a sex-difference. This approach shows that in most cases, Bayes Factors for models considering sex versus the ones not considering sex showed that assuming no effect of sex was about twice as good at predicting the data than models assuming an effect of sex. This means that overall we certainly do not have evidence for an effect of sex, but that we also do not have strong evidence for the absence of an effect of sex. We thus summarize our data as showing that emotional contagion is roughly similar across the sexes in the rats. A larger group size would be necessary to rigorously exclude the presence of even small effects.

In our paradigm, we also quantified how much the demonstrator’s freezing is influenced back by the observer’s freezing. Consistent with our previous work, which has shown that taking the observer’s freezing into account helps predict how much a demonstrator freezes in response to shock ^16,29^, we find in both our regression and Granger causality approach, evidence that the observer’s freezing influences the demonstrator’s freezing. This phenomenon, traditionally called social buffering ^44^, was also not sex-dependent in our sample.

In contrast to the lack of a significant sex effect on emotional contagion, we do find a robust sex difference in demonstrator’s freezing, with females freezing half as much as males in response to shocks, a finding confirmed in a second group (Fig. 4B). This difference then feeds onto the observers, which also show, an albeit smaller, sex difference, which can, however, be entirely explained by the sex difference in demonstrator freezing. Intriguingly, we do not find this sex difference during the pre-exposure session, where males and females also received electroshocks. Several differences exist between what observer animals experience during pre-exposure and what demonstrators experience during the emotional contagion test. First, the parameters of the footshocks differ (Fig. 4). Observers during pre-exposure experienced 4 footshocks, 0.8mA, 1 sec long, 200-260sec random inter-shock interval while demonstrator received 5 footshocks: 1.5mA, 1 sec long, 120 or 180sec inter-shock interval. However, we also found no sex-difference in the pre-exposure but a sex-difference during the emotional contagion test when only testing the first 110s of the first 3 shocks, suggesting that the difference in the number of footshocks and the intershock interval are perhaps unlikely to explain the difference in sex-effects. Shock intensity, however, could play an important role. Second, in pre-exposure animals encounter the pre-exposure context for the first time on the day they receive shocks, while demonstrators in the test context have been habituated for three days to that context. Finally, animals during pre-exposure are alone, while demonstrators in the test situation are paired with their cage-mate. Whether the strong sex difference in shock-triggered freezing we observe in our test situation is specific to the shock intensity, level of habituation and/or social buffering remains for future experiments to test.

Our study also has a number of important limitations. First, we did not test pairs of mixed sexes, with a female demonstrator paired with a male demonstrator and vice versa. Future experiments should explore if emotional contagion might be stronger within a gender than between genders. Second, we have only explored a small number of animals in our study, and small sex-differences may thus have evaded our analysis. Third, we have only explored emotional contagion in rats that have been pre-exposed to shocks. We have previously shown that pre-exposure of observers to shocks does increases the amount of freezing in observers of both sexes quite dramatically ^16,29^. Whether there might be sex-differences in the social transmission of shock-naïve observers might be worth exploring in the future. Finally, we have only explored the sex-difference in Long-Evans rats. In the future, exploring this effect in other rat strains and in mice would be exciting.

In summary, we show that although we find significant sex differences in how demonstrators react to a shock, the emotional contagion process, that transmits this reaction across individuals does appear to be sex-independent in Long Evan rats. This echoes two observations we have recently made regarding familiarity in the same paradigm. First, we observed that emotional contagion is independent of how long male observers and demonstrators have been housed together ^29^. Second, we have compared emotional contagion across male Long-Evans and Sprague-Dawley rats. We found that akin to female Long-Evans, Sprague-Dawley male demonstrator rats freeze significantly less to the electroshock. However, the degree of social transmission, as estimated using the slope of a regression or Granger causality, did not depend on the strain. Taken together, this shows that emotional contagion in our paradigm is similar across different sexes, different strains and different levels of familiarity. This finding is compatible with the notion that emotional contagion primarily serves a purpose similar to Eaves-dropping across animals ^45–47^, namely the social detection of danger ^29^. If a rat witnesses another rat express distress, this is a valuable, selfish danger signal that the recipient can use as an indicator of danger that should trigger freezing. In this selfish, danger-detection view, one would not necessarily expect that females should show more emotional contagion than males.

Overall, the lack of behavioral sex-effect on emotional contagion we find here thus does not lead us to question whether the neuroscience work done to uncover the neural basis of emotional contagion in male rodents ^21,26,28,30,31^ also applies to females. However, similar behavioral outcomes could arise from slightly different neural circuits across genders, warranting a similar examination of sex-differences also in markers of neural activity.

Finally, it is important, to recognize that there might be a fundamental difference between emotional contagion, as measured using the social transmission of distress and freezing, and prosocial motivation, as measured using directed helping. The latter, as measured using the latency to liberate a trapped conspecific, is stronger in more familiar animals and in females ^43,48^. The exact causal relationship between these two phenomena remains to be explored.

## 4. Methods

### Subjects

Ten male and nine female Long Evans rats (observers) were bred in house at the animal facility of the Netherlands Institute for Neuroscience. We bred these animals inhouse as part of a larger study exploring early life stress using a limited bedding and nesting manipulation ^replication of 49^ but we mainly present the data from the control condition here (except for Fig. 4). The breading led to slightly more males, explaining the slight difference in numbers. Upon weaning animals were housed in same-sex groups of 4, maintained at ambient room temperature (22-24°C, 55% relative humidity, SPF, type IV cages, on a 12:12 light-dark cycle: lights on at 07:00) till 6 weeks of age. As in previous studies, we always ordered demonstrators from Janvier ^17,28,29^, here as well, we ordered ten male and nine female Long Evans rats (demonstrators; 6 weeks of age) from Janvier Labs (France). Upon arrival, animals were pair-housed with observers (same-sex pairs) in type III cages with wooden block toys, on a reversed 12:12 light-dark cycle (lights off at 07:00). Food and water were provided *ad libitum*.

### Ethics Statement

In compliance with Dutch and European law and institutional regulations, all experimental procedures were preapproved by the Centrale Commissie Dierproeven of the Netherlands (AVD801002015105) and by the welfare body of the Netherlands Institute for Neuroscience (IVD, protocol number NIN161108) and by the welfare body of the Netherlands Institute for Neuroscience (IVD, protocol number NIN161108) in accordance with the Experiments on Animals Act (WOD) with its amendment on 18 December 2014 and EU directive 2010/63/EU. All animals from this experiment were handled at least once a week prior to the experimental start and habituated to the experimental room and setup to reduce unnecessary stress. The welfare of the animals was monitored throughout, and no animals had to be sacrificed because of signs of illness. After experiments, the animals were euthanized by CO_2_ inhalation, starting with 40% O_2_ mixed with 60% CO_2_ until animals were in deep sleep (as checked by the rear reflex response and breathing depth and frequency) and then switched to 100% CO_2_ for at least 15 minutes until no breathing or heartbeats were detected.

### Setup

All tests were conducted in a two-chamber apparatus (each chamber L: 24cm x W: 25m x H: 34cm, Med Associates, Inc.). Each chamber consisted of transparent Plexiglas walls and stainless-steel grid rods. The compartments were divided by a transparent perforated Plexiglas separation, which allowed animals in both chambers to see, smell, touch and hear each other. For shock pre-exposure and the emotional contagion tests, one of the chambers was electrically connected to a stimulus scrambler (ENV-414S, Med Associates Inc.). For video recording of the rats’ behaviors, a Basler GigE camera (acA1300-60gm) was mounted on top of the apparatus controlled by Media Recorder (Noldus, the Netherlands).

### Experimental procedures

The experimental procedures consisted of habituation, pre-exposure and test phases (Fig. 1). Ten days prior to the emotional contagion test, all animals were handled every other day for 3 minutes/day. To habituate animals to the testing conditions, animal dyads were transported and placed in the testing apparatus for 20 mins per day for three sessions. The testing apparatus was cleaned with lemon-scented dishwashing soap and 70% alcohol between each dyad. To enhance the emotional contagion response to the distress of the demonstrators, observer animals experienced a shock pre-exposure session^16,29^. The shock pre-exposure was conducted in one of the chambers of the test apparatus. To prevent contextual fear on the test day, the walls of the chamber were coated with black and white striped paper, the background music was turned off, the apparatus was illuminated with bright white light and the chamber was cleaned with rose-scented dishwashing soap and vanilla aroma drops for the pre-exposure session. Observers were individually placed in the apparatus and after a 10-minute baseline plus a random interval (∼230sec), four footshocks (each: 0.8mA, 1 sec long, 200-260sec random inter-shock interval) were delivered. After the shock pre-exposure session, animals were placed for 1 hour in a neutral cage prior to returning to their home cage. The emotional contagion test setup was illuminated with dim red light, cleaned using a lemon-scented dishwashing soap followed by 70% alcohol, and background radio music was turned on. Observer-demonstrator dyads were transported during their dark-cycle to the testing room and animals were placed in the corresponding chamber of the testing apparatus. For all dyads, following a 12-minute baseline, the demonstrators experienced five footshocks (each: 1.5mA, 1 sec long, 120 or 180sec inter-shock interval). Following the last shock, dyads were left in the apparatus for 2 additional minutes prior to return to their home cage.

### Behavior scoring

The behaviors of observers and demonstrators during the emotional contagion test and pre-exposure were manually scored by 2 experienced researchers and using the open-source Behavioral Observation Research Interactive Software (BORIS, Friard & Gamba, 2016). Freezing, defined as lack of movement except for breathing movements, was continuously scored throughout the baseline and the shock period. Freezing had to last for at least 2 seconds to be scored as freezing. To create a continuous time series, freezing moments extracted from the Boris result files were recoded as 1 and non-freezing moments as 0 using Matlab (MathWorks inc., USA). We also scored rearing/climbing behaviors (i.e. front paws/both front and rear paws away from the floor and not grooming) using Boris.

### Statistics

For sections 2.1 and 2.2, freezing time was calculated as the sum of all freezing moments in a certain epoch and freezing percentage was calculated as the total freezing time divided by the total time of the epoch. Baseline period (1st epoch) was defined as the first 720 seconds of the emotional contagion test and the shock period (2nd epoch) was defined as the 720 seconds following the first shock (approx. 720 seconds from the start of the test). Statistics were computed using JASP (version 0.11.2, https://jasp-stats.org/). Bayesian results are reported as Bayes Factors, with the index specifying whether it is the Bayes Factor in favor of the hypothesized effect *H*_1_ or the null hypothesis *H*_0_, with BF_10_ P(data|*H*_1_)/P(data|*H*_0_) and BF_01_= P(data|*H*_0_)/P(data|*H*_1_) BF_10_>3 represents moderate evidence for an effect, BF_10_<1/3 for the absence of an effect, and the reverse is true for BF_01_. BF values around 1 indicate the data is similarly likely under *H*_0_ and *H*_1_, and cannot adjudicate in favor of either. For model comparisons, BF_incl_ represents the ratio between the likelihood of models including, divided by those excluding a particular factor. BF_incl_>3 is interpreted as moderate evidence that the inclusion of this particular factor improves the model. BF_incl_<1/3 is interpreted as moderate evidence that this particular factor does not improve the model). Default priors are used throughout. This includes for ANCOVAs: *r*=0.5 for the effect of sex (a fixed effect between-subject factor) and *r*=0.354 for covariates; and for comparisons between groups a Cauchy with a scale of 0.707.

### Granger causality

Granger causality is a statistical concept of causality that is based on prediction ^50^. If a signal *X*1 “Granger-causes” (or “G-causes”) a signal *X*2, then past values of *X*1 should contain information that helps predict *X*2 above and beyond the information contained in past values of *X*2 alone. In this study, *X*1 and *X*2 were binary time series of freezing of the demonstrator and freezing of the observer (freezing coded as 1 and not-freezing coded as 0) on a second-to-second basis. The freezing of the observer at a certain time point (*X2(t)*) can be estimated either by its own history plus a prediction error (reduced model, 1) or also including the history of the freezing of the demonstrator (full model, 2).

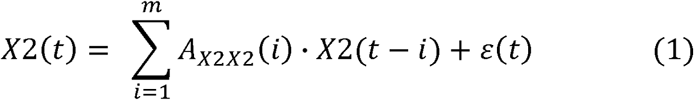

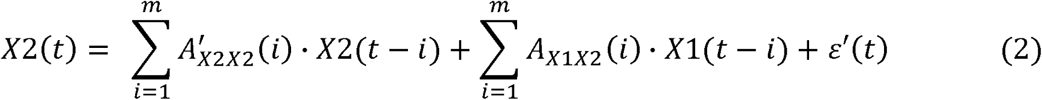

In equations 1 and 2, *t* indicates the different time points (in steps of 1s), *A* represents the regression coefficients and *m* refers to the model order which is the length of the history included. Granger causality from the freezing of the demonstrator to the freezing of the observer (i.e. *X*1→*X*2) is estimated by comparing the full model (2) to the reduced model (1). Mathematically, the log-likelihood of the two models (i.e. G-causality value *F*) is calculated as the natural logarithm of the ratio of the residual covariance matrices of the two models (3).

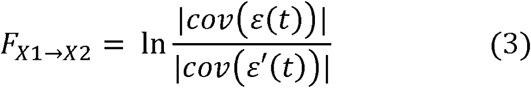

This G-causality magnitude has a natural interpretation in terms of information-theoretic bits-per-unit-time ^51^. In this study, for example, when G-causality from the demonstrator to the observer reaches significance, it indicates that the demonstrator’s freezing can predict the observer’s freezing and that there is information flow from the demonstrator to the observer. Jumping responses of the demonstrator to the foot shocks were also taken into account and a binary time series of this behavior was included as *X*3 (jumping coded as 1 and not-jumping coded as 0).

The algorithms of the Multivariate Granger Causality (MVGC) Toolbox ^51^ in MATLAB were used to estimate the magnitude of the G-causality values. First, the freezing time series of the demonstrators and the observers were smoothed with a Gaussian filter (size = 300s, sigma =1.5). The MVGC toolbox confirmed that each time series passed the stationary assumption for Granger causality analysis. Then, the optimal model order (*m*, the length of history included) was determined by the Akaike information criterion (AIC) for the model including all observer-demonstrator dyads. The optimal model order is a balance between maximizing goodness of fit and minimizing the number of coefficients (length of the time series) being estimated. For this dataset, the model order of 19 was estimated to be the best fit for the model including all dyads and thus it was fixed at 19 for the subsequent dyad-wise analysis. To test the differences of the G-causality values sexes, we used the g-causality values obtained from analysis only considering the freezing during the shock epoch of a given pair of rats, and compared them across groups using SPSS and JASP.

## Acknowledgments

We thank Nine Kompier for her help with scoring the freezing behavior of the animals.

## Data and materials availability

All data (except for the movies from the behavior) will be uploaded on Mendeley Data before publication. Until then, it can be downloaded from https://www.dropbox.com/sh/3s65glzdvbmdchg/AACorwOSO9Q7t3E8WkKGyXkNa?dl=0.

